# A Comprehensive Assessment of the Genetic Determinants in *Salmonella* Typhimurium for Resistance to Hydrogen Peroxide Using Proteogenomics

**DOI:** 10.1101/115360

**Authors:** Sardar Karash, Rohana Liyanage, Abdullah Qassab, Jackson O. Lay, Young Min Kwon

## Abstract

*Salmonella* is an intracellular pathogen that infects a wide range of hosts and can survive in macrophages. An essential mechanism uses by the macrophages to eradicate *Salmonella* is production of reactive oxygen species. Here, we used proteogenomics to determine the candidate genes and proteins that have a role in resistance of *S.* Typhimurium to H_2_O_2_. For Tn-seq, a highly saturated Tn5 insertion library was grown *in vitro* under either 2.5 (H_2_O_2_L) or 3.5 mM H_2_O_2_ (H_2_O_2_H). We identified two sets of overlapping genes that are required for resistance of *S*. Typhimurium to H_2_O_2_L and H_2_O_2_H, and the results were validated via phenotypic evaluation of 50 selected mutants. The enriched pathways for resistance to H_2_O_2_ included DNA repair, aromatic amino acid biosynthesis (*aroBK*), Fe-S cluster biosynthesis, iron homeostasis and a putative iron transporter system (*ybbKLM*), flagellar genes (*fliBC*), H_2_O_2_ scavenging enzymes, and DNA adenine methylase. Proteomics revealed that the majority of essential proteins, including ribosomal proteins, were downregulated upon exposure to H_2_O_2_. A subset of proteins identified by Tn-seq were analyzed by targeted proteomics, and 70% of them were upregulated upon exposure to H_2_O_2_. The identified candidate genes will deepen our understanding about mechanisms of *S*. Typhimurium survival in macrophages, and can be exploited to develop new antimicrobial drugs.

## Introduction

*Salmonella* is a Gram-negative bacterium that infects humans and animals. *Salmonella enterica* has numerous serovars, which include typhoidal and non-typhoidal strains. In contrast to the typhoidal salmonellae which are human restricted pathogens, the non-typhoidal salmonellae (NTS), serovar Enteritidis and Typhimurium, are able to infect a wide range of hosts, causing gastroenteritis^1^. The NTS strains, including *Salmonella enterica* serovar Typhimurium, account for 11% (1.2 million cases) of the total foodborne illnesses caused by different pathogens in the United States^2^. It has been estimated that *Salmonella* is responsible for 93.8 million cases of gastroenteritis, leading to 155,000 deaths worldwide annually^3^. The pathogen remains a continuous threat to the food safety, and public health.

To initiate an infection and survive inside the host, *Salmonella* needs to overcome a myriad of host defense mechanisms. As *Salmonella* reaches the intestine and breaches the epithelial tissue, it enters the macrophages and activates different virulence strategies in order to survive and replicate in them^4^. An essential mechanism uses by the phagocytes to kill and eradicate *Salmonella* is production of reactive oxygen species (ROS). Hydrogen peroxide (H_2_O_2_), superoxide anion 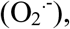 and the hydroxyl radical (HO.) are derivatives of ROS. The short-lived 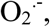 produced by the NADPH-dependent phagocytic oxidase, quickly dismutates into H_2_O_2_, which diffuses across semipermeable bacterial cell membranes. Eventually, Fe^2+^ reduces H_2_O_2_ to HO. via the so called Fenton Reaction^5–7^. The ROS, including H_2_O_2_, can damage DNA, iron-sulfur cluster-containing proteins, and other biological molecules in the bacterial cells^8–10^.

Numerous genetic factors and proteins that are important for resistance of *S.* Typhimurium to H_2_O_2_ have been discovered and the underlying mechanisms have been explored^11, 12^. A various approaches and techniques have been employed to study global response of *Salmonella* or related bacteria to H_2_O_2_ *in vitro* as a model system to simulate the bacterium’s response to ROS in phagocytic cells: (i) Two-dimensional gel electrophoresis identified H_2_O_2-_induced proteins in *Salmonella*^13^, (ii) DNA microarray identified H_2_O_2_ induced genes in *E. coli*^14^, and (iii) RNA-seq identified H_2_O_2_ induced genes in *Salmonella*^15^. Yet, the factors required for fitness under the given condition cannot be identified with high confidence based on the analysis of transcriptomics or proteomics data^16^. Microarray-based tracking of random transposon insertions was used to identify numerous genes in *Salmonella* that are required for survival in mice and macrophages^17–18^. However, the genetic factors responsible for resistance to ROS cannot be sorted out among all of the genetic factors identified in the study that are required for fitness in the presence of multiple host stressors.

To shed more insights into the underlying mechanisms of *Salmonella* resistance to H_2_O_2_, more direct approach linking the gene-phenotype relationships in a genome-wide scale would be necessary. Tn-seq is a powerful approach to allow direct and accurate assessment of the fitness requirement of each gene on the entire genome of a prokaryotic organism^19^. In Tn-seq method, a saturated transposon insertion library (input) is exposed to a selective condition, and the mutant population altered through the selection (output) is recovered. Then, the genomic junctions of the transposon insertions are specifically amplified and sequenced from both input and output pools by high-throughput sequencing. The gene fitness can be obtained by calculating the change in relative abundance of the sequence reads corresponding to each gene in the entire genome between the two pools. Tn-seq has been employed to assign gene functions to *Salmonella* genomes in numerous studies: (i) Previously, our lab identified conditionally essential genes that are required for growth in the presence of bile, limited nutrients, and high temperature^20^, (ii) The genes required for intestinal colonization were identified in chickens, pigs, and cattle^21^, (iii) Candidate essential genes and genes contributing toward bile resistance were identified^22^, (iv) Core conserved genes for growth in rich media were identified in serovars Typhi and Typhimurium^23^. In addition to Tn-seq, electrospray ionization liquid chromatography tandem mass spectrometry (ESI-LC-MS/MS) is a powerful approach for identifying and quantifying proteins in a large scale. The system-wide protein regulation can be determined using mass spectrometry signal intensities of tryptic peptides obtained from two different culture conditions^24^. The post-translational modification in proteins can be revealed by using proteomic analysis^25^. Many studies took advantage of proteomic analysis of *Salmonella*. However, to the best of our knowledge, this study is the first to investigate proteogenomics of a bacterium by combining Tn-seq and proteome analysis simultaneously to the same stressor.

In this work, we used Tn-seq method and proteomic analysis in combination to determine system-wide responses of *S.* Typhimurium to two different concentrations of H_2_O_2_ (H_2_O_2_L and H_2_O_2_H). We obtained a comprehensive list of 137 genes that are putatively required for the resistance of *S.* Typhimurium 14028 to H_2_O_2_. The role of 50 selected genes in resistance to H_2_O_2_ were determined by phenotypic evaluation of the individual deletion mutants. Also, we identified a set of 246 proteins that are differentially expressed in response to H_2_O_2_, using data-dependent acquisition (DDA) proteomics, which are largely overlapped with the genes identified by Tn-seq; targeted proteomics showed 70% of the proteins identified by Tn-seq were upregulated by H_2_O_2_. In addition to the genes of *S.* Typhimurium previously known to be important for resistance to H_2_O_2_, we identified approximately 80 genes that have not been previously associated with resistance to oxidative stress. The results of this study highlighted that the genes in aromatic amino acid biosynthesis, *aroB* and *aroK*, and iron homeostasis, *ybbK*, *ybbL*, and *ybbM*, are crucially important for growth fitness under H_2_O_2_ stress. The identified candidate genes will expand our understanding on the molecular mechanisms of *Salmonella* survival in macrophages, and serve as new antimicrobial drug targets.

## Results and Discussion

### The H_2_O_2_ concentrations and the selections of Tn5 library

First, we sought to determine the growth response of wild type *S.* Typhimurium 14028 cells in LB media containing varying concentrations of H_2_O_2_. The wild type cells were grown in LB media that contain different concentrations of H_2_O_2_ in 96-well plates. After evaluating the growth rates for the cultures, 2.5 and 3.5 mM H_2_O_2_ were chosen for Tn-seq selections in our study, and termed H_2_O_2_L and H_2_O_2_H, respectively. In comparison to *Salmonella* grown in LB media with no H_2_O_2_, H_2_O_2_L and H_2_O_2_H reduced the growth rates by 10% and 28%, respectively (Fig 1A). The lag time increased by a 5.7-fold (0.5 vs. 2.9 hr), and an 11-fold (0.5 vs. 5.6 hr) in H_2_O_2_L and H_2_O_2_H, respectively. The maximum OD_600_ decreased by only 1% for the H_2_O_2_L and 2% for the H_2_O_2_H in comparison to LB media (Fig 1A).

**Figure 1.**
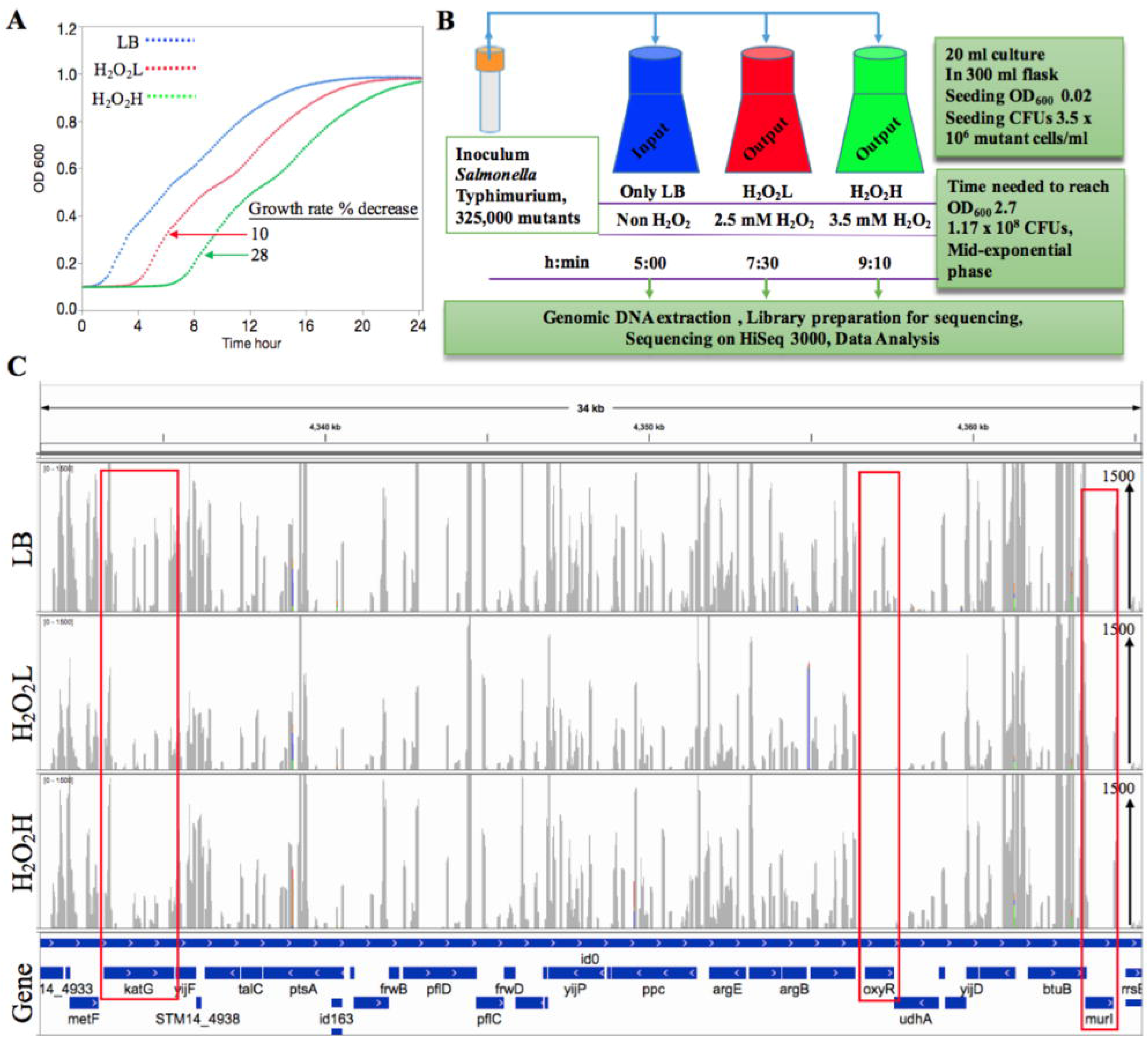
Study design and identification of the genes required for H_2_O_2_ resistance using Tn-seq. (A) The effect of H_2_O_2_ on the growth rate of wild type *Salmonella* Typhimurium. An overnight culture of bacteria was diluted 1:200 in the LB medium contains either 2.5 mM H_2_O_2_ (H_2_O_2_L), 3.5 mM H_2_O_2_ (H_2_O_2_H), or LB without H_2_O_2_ was used as control. The cultures were incubated at 37°C for 24 h in a 96-well plate. The reduced growth rates for the H_2_O_2_ were in comparison to the control. In the all growth curve figures in this work, the blue color represents LB (no H_2_O_2_), the red is H_2_O_2_L, and green is H_2_O_2_H. (B) Schematic representation of the Tn-seq study. The *Salmonella* transposon mutant library was inoculated into LB and LB contains H_2_O_2_L or H_2_O_2_H. The three cultures were grown until they reached mid-exponential phase. The DNA was extracted from each culture and subjected to library preparation, sequencing, and data analysis. (C) The Tn-seq profile of the three conditions. It shows 34 kb of the *Salmonella* genome which starts with *metF* gene and ends with *rrsB*, horizontal axis. The height of vertical axis represents number of reads which is 1500 sequencing reads. The highlighted genes in red are *katG*, catalase peroxidase, was tolerated insertions in the both H_2_O_2_ conditions; *oxyR* was not tolerated insertions in presence of H_2_O_2_ and indicated that the gene is required to H_2_O_2_ resistance; *murI*, glutamate racemase, was not tolerated any insertions at all and it was considered an essential gene, *murI* is required for the biosynthesis of a component of cell wall peptidoglycan.

For the selection of Tn5 library, 20 ml cultures in 300 ml Erlenmeyer flasks containing LB, H_2_O_2_L, or H_2_O_2_H were inoculated with the same Tn5 library at the seeding CFUs of the library at 3.5×10^6^. This seeding level provided ∼10 CFUs for each Tn5 insertion mutant in the library. The cultures were grown until the mid-exponential phase, in which the CFUs reached 1.17×10^8^ (SE 0.01×10^8^). It required 7.5 and 9.2 h to reach the cell density as measured by optical density for H_2_O_2_L and H_2_O_2_H, respectively, in contrast to 5 h for LB medium (Fig 1B). We observed some differences in growth responses between the cultures in a 96-well plate and in a 300-ml Erlenmeyer flask. The optical density readings by the plate reader was different in comparison to those by Bio-photometer that we used to measure optical density of the culture in the flask. As a result, the growth curve in Fig 1A which was based on 96-well plate reader, dose not match exactly with the time required for the Tn5 library to reach the target mid-exponential phase in the flask cultures. In addition, we observed that the H_2_O_2_ is stable in LB media free of *Salmonella* during the window of time used for the library selection process (Fig. S1), which was also supported by Bogomolnaya et al.^26^.

### Preparation of Tn5-seq amplicon library

The *Salmonella* mutants were generated by using the delivery plasmid pBAM1 via conjugation. A total of 325,000 mutant colonies were recovered from 50 plates. Each mutant contained a single random insertion of Tn5 transposon in the chromosome or plasmid according to DNA sequencing of Tn5-junction sequences for a small set (n = 71) of randomly selected Tn5 mutants. We found a significant portion (∼20%) of the mutants in the library that were not genuine Tn5 insertions, but the mutants generated as a result of pBAM1 integration into chromosome as determined by their ability to grow in the presence of ampicillin. To prevent the Illumina sequencing reads of being wasted on sequencing Tn5 junctions from these cointegrants, we digested genomic DNA of the input and output libraries with PvuII, which digests immediately outside the inverted repeats on both sides of Tn5. The digested DNA was then used to prepare Tn-seq amplicon library as described in Materials and Methods. Our Tn-seq data analysis indicated that our strategy of removing the DNA sequences originating from cointegrants was effective because only 0.55% of the total HiSeq reads corresponding to Tn5-junctions matched to pBAM1. It should be possible to remove them completely by ensuring complete digestion of genomic DNA with PvuII. The method for Tn-seq amplicon library we developed and used in this study has multiple advantages over other Tn-seq protocols, because our method requires only 100 ng of the genomic DNA, and the whole process can be completed in a day^27^. When the extension step in the protocol was performed using a conventional 20 nucleotide primer, and the final products of exponential PCR were separated on agarose gel electrophoresis, even the negative controls (the wild type genomic DNA or mutant library genomic DNA without linear extension) showed smear patterns of nonspecific background amplification. However, when dual priming oligonucleotide (DPO) primer was used in place of the conventional primer for linear extension, non-specific background amplification was completely disappeared. Therefore, we adopted the DPO primer in linear extension step for all library samples in this study. Then, the single-stranded extension products were C-tailed, and used as templates for the exponential PCR step using nested primer specific to Tn5 and poly G primer that contain Illumina adapter sequences along with sample index sequences (Fig. S2). The final PCR products were separated on an agarose gel, and the fragments within the range of 325-625 bp were gel-purified. After pooling of multiple samples, the combined library was sequenced on a HiSeq 3000.

### Summary of Tn-seq DNA analysis

After de-multiplexing and C-tail trimming of all sequence reads, ∼72 million reads of Tn5-junctions with mean read length of 94 bp were obtained. The number of the reads mapped to the complete genome of *S.* Typhimurium 14028 were ∼25, 15, and 19 million for LB, H_2_O_2_L, and H_2_O_2_H, respectively. The number of unique insertions on the chromosome were 125,449 in the input library, excluding the plasmid (Table S1). On average, Tn5 was inserted in every 39 bp. Number of raw reads per open reading frame (ORF) for H_2_O_2_L was plotted over the corresponding number of H_2_O_2_H, which yielded an R^2^ of 0.91, indicating the mutants in the input library quantitatively responded in a similar way for both H_2_O_2_L and H_2_O_2_H as expected (Fig. S3). The insertions were mapped to 5,428 genes or 8,022 genes/intergenic regions. Interestingly, the ORF STM14_5121, which is 16.7 kbp long, had the highest number of insertions (∼700 insertions) and reads (0.25 M).

### Comparison of various bioinformatics pipelines for Tn-seq data analysis

We used 3 different Tn-seq analysis tools to identify the genes and compare the results across the methods with the goal of comprehensive identification of “all” genes required for resistance to H_2_O_2_. The first tool, ARTIST^28^, created small non-overlapping genomic windows of 100 bp and the reads from each window were arbitrarily assigned into the middle of the window. The default normalization script of the tool was used. Then, the relative proportions of insertion sites in the output library versus the input were tabulated. Mann-Whiney *U* (MWU) test was used to assess the essentiality of the locus. To consider a gene/intergenic region conditionally essential for growth in the presence of H_2_O_2_, *p* value had to be ≤ 0.05 in 90 of the 100 conducted MWU tests. Subsequently, 20 genes and 1 intergenic region were identified for H_2_O_2_L and 4 genes for H_2_O_2_H (Table S2). We speculate the reason that more genes were identified for H_2_O_2_L in comparison to H_2_O_2_H, was partially due to the lower number of total reads of H_2_O_2_L as compared to H_2_O_2_H, even though the read numbers of H_2_O_2_L was normalized to those of the input.

The second tool, Tn-seq Explorer^29^, counted insertions in overlapping windows of a fixed size. Using a 550 bp window size, each annotated gene was assigned an essentiality index (EI) which is determined mainly based on the insertion count in a window in this gene. The bimodal distribution of insertion counts per window divided the essential genes to the left and the non-essential genes to the right. To find conditional essential genes, the EI of the output was subtracted from the EI of the input. The genes with negative ΔEI were ranked based on the change in read fold change (Log2 (H_2_O_2_L or H_2_O_2_H/Input)). We found 114 consensus genes between H_2_O_2_L and H_2_O_2_H that had at least four-fold reduction in H_2_O_2_H read counts as compared to the input. The four-fold reduction (Log2FC = −2) threshold was chosen based on our validation study of Tn-seq data by single mutant assays (Table S2).

The third tool, TRANSIT^30^, determined read counts of genes in the input and output library. The differences of total read counts between the input and outputs were obtained. The insertion sites were permutated for a number that is specified by the user (we used 10,000 sample). This sampling for each gene gave difference in read counts. The *p* value was calculated from the null distribution of the difference in read counts. We identified 8 and 21 genes for the H_2_O_2_L and H_2_O_2_H, respectively, using a *p* value ≤ 0.05 (Table S2).

The combined list of the genes identified by the 3 Tn-seq analysis tools for both H_2_O_2_L and H_2_O_2_H included 137 genes (Table S2). All of the genes on this list are expected to have a role in conferring resistance to H_2_O_2_ and allow *Salmonella* to survive and replicate in the presence of H_2_O_2_ *in vitro*. Of the 21 genes identified by TRANSIT, 19 of these genes were also identified by Tn-seq Explorer, but only 3 out of this 21 were identified by ARTIST. The 19 genes were *hscA*, *rbsR*, *fepD*, *efp*, *oxyR*, *polA*, *ybaD*, *aroD*, *ruvA*, *xthA*, *dps*, *aroB*, *uvrD*, *tonB*, *uvrA*, *aroK*, *ybbM*, *lon*, and *proC*. Two genes, *fepD* and *xthA*, were identified by the all 3 methods and for both conditions.

The 3 Tn-seq analysis tools are very valuable for Tn5 data analysis, but each tool has its own advantages and disadvantages. For ARTIST, (i) the user must know how to run scripts in Matlab software, (ii) the analysis is very slow on a personal computer with the HiSeq data, (iii) it has only one method for normalization, but (iv) it can search for essentiality in the intergenic regions. For Tn-seq Explorer, (i) there is no data normalization, and (ii) prediction on small genes is prone to be inaccurate, but (iii) its very user-friendly and runs fast. For TRANSIT, (i) the user should have some knowledge on running scripts on terminal, (ii) it may need some modification in its Python script according to the way the library was prepared for sequencing, and (iii) a few software packages should be installed on the computer as TRANSIT pre-requisites, but (iv) it does have 6 different methods for data normalization and it runs very fast on a personal computer. Although ARTIST and Tn-seq Explorer are very useful tools for Tn-seq data analysis, we prefer using TRANSIT in our future data analysis for conditionally essential genes. In the following sections, we continued the downstream analysis mainly based on the 137 genes that include all of the genes identified by all 3 methods.

### The enriched pathways for resistance to H_2_O_2_

In order to categorize the identified genes that are required for *Salmonella* resistance to the H_2_O_2_, the 137 genes were subjected to pathway enrichment analysis using DAVID Bioinformatics Resources 6.7, NIAID/NIH^31^. A total of 15 KEGG pathways^32^ were recognized for 69 genes on the list. The enriched pathways include homologous recombination (*ruvC*, *polA*, *ruvA*, *ruvB*, *priB*, *recA*, *recR*, *holC*, *holD*, *recC*, *recG*), nucleotide excision repair (*uvrD*, *polA*, *uvrA*, *uvrC*), mismatch repair (*dam*, *uvrD*, *holC*, *holD*), RNA degradation (*pnp*, *hfq*, *ygdP*), purine and pyrimidine metabolism (*apaH*, *polA*, *pnp*, *arcC*, *spoT*, *holC*, *holD*, *cmk*, *dcd*, *pnp*), phenylalanine, tyrosine and tryptophan biosynthesis (*aroD*, *aroB*, aroA, *aroK*, *aroE_2*), arginine and proline metabolism (*proC*, *arcC*), glycolysis and gluconeogenesis (*crr*, *pgm*, *tpiA*), oxidative phosphorylation (*atpG*, *atpA*, *cydA*), DNA replication (*polA*, *holC*, *holD*), and flagellar assembly (*fliJ*, *fliD*, *flhD*, *fliC*). Since KEGG was not able to recognize many genes on the list, we used SP_PIR_Keywords of functional categories, which recognized majority of the genes and categorized them into 55 functional categories (Table S3), excluding 15 uncharacterized genes (ORFs). Among these categories were stress response (*rpoE*, *lon*, *dnaJ*, *hfq*, *yaiB*), iron (*dps*, *entD*, *iscA*, *yjeB*, *yhgI*), and transcription regulation (*rcsA*, *oxyR*, *rpoE*, *yjeB*, *arcA*, *argR*, *rbsR*, *rpoS*, *fadR*, *rcsB*, *furR*, *flhD*).

### Validation of Tn-seq results using individual mutants

For the selected 50 genes among the 137 genes identified by Tn-seq, the growth phenotype was determined using individual single deletion mutants in LB, H_2_O_2_L, and H_2_O_2_H. The genes were considered to play a role in resistance to H_2_O_2,_ if (i) lag phase time increased, (ii) growth rate reduced or (iii) maximum OD_600_ decreased in the presence of H_2_O_2_ in comparison to the wild type strain grown in the same conditions. Of the 50 single deletion mutants, 42 mutants were shown to have a role in resistance to H_2_O_2_ (Fig. 2 and Table S4). One gene, *yhaD*, was identified by all 3 analysis tools, but it did not show the expected phenotype. The *fliD* was also identified by ARTIST, but did not show any phenotype distinguishable from the wild type. The remaining 6 genes that did not show the phenotype was identified by Tn-seq Explorer. Based on the results of the individual mutant assay, we conclude that 84% (42/50) of the genes identified by the Tn-seq analysis and tested using single deletion mutants have a role for resistance to H_2_O_2_. These results indicate that our Tn-seq analysis identified the genes in *S.* Typhimurium that are required for the wild type level resistance to H_2_O_2_ with high accuracy.

**Figure 2.**
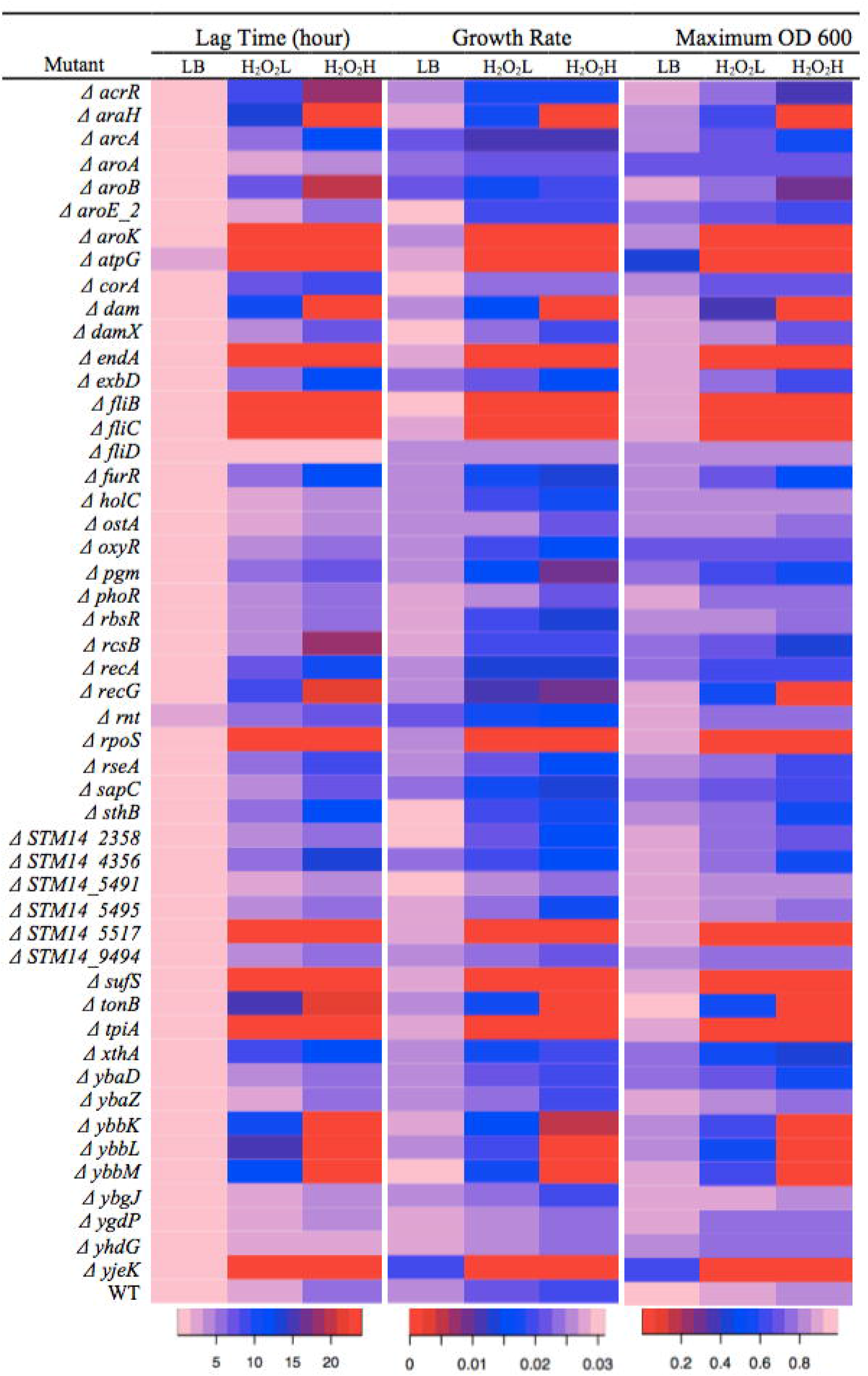
Growth curve of 50 mutants and a wild type *Salmonella* in LB, H_2_O_2_L and H_2_O_2_H. The lag phase time, growth rate, and maximum OD_600_ of the individual *Salmonella* Typhimurium mutants and the wild type in the growth conditions of LB (no H_2_O_2_), H_2_O_2_L and H_2_O_2_H. The overnight cultures of the mutants and wild type *Salmonella* were diluted 1:200 in the LB medium, and the LB contains either 2.5 mM H_2_O_2_ (H_2_O_2_L) or 3.5 mM H_2_O_2_ (H_2_O_2_H). The cultures were incubated at 37°C for 24 h in a 96-well plate and the OD_600_ was recorded every 10 min. The lag phase time, growth rate, and maximum OD_600_ were calculated and shown here as a graphical representation. The pale pink color indicates a short lag phase time, a high growth rate, and a high OD_600_. The red color indicates that the bacteria was stayed in a lag phase, growth rate was close to a zero, and the OD_600_ of the culture was not raised in the 24 h time of assays. The data of this figure can be found in Table S4.

### Proteomics of H_2_O_2_ response

With ESI-LC-MS/MS in data-dependent acquisition (DDA) mode, the protein regulation was determined using MS1 filtering technique that skyline software offers^33^. It uses signal intensities of tryptic peptides derived from the proteins of wild type strain grown in the presence of H_2_O_2_ in comparison to the control (LB). As described in Materials and Methods section, trypsin digestion of the protein extracts under different conditions generates tryptic peptides that are uniquely related to individual proteins. Tryptic peptides separated by liquid chromatography from the complex samples were first subjected to simple mass measurement (MS1) followed by intensity dependent fragmentation of these peptide ions to produce sequence specific fragment ions by collision-induced dissociation (MS/MS). Tryptic peptides were then identified using these sequence specific fragment ions via MASCOT database search software^34^, where the sequence specific fragment ions were matched to the proteins in *S.* Typhimurium 14028S reference proteome database^24, 35^. This method of protein analysis is normally referred to as data dependent analysis (DDA). At the beginning of data analysis, the H_2_O_2_L and H_2_O_2_H data were compared to LB separately, however it turned out that comparison was not sensitive enough to differentiate between H_2_O_2_L and H_2_O_2_H conditions. Hence, the data of H_2_O_2_L and H_2_O_2_H were combined for analysis in comparison to LB. We identified 1,104 proteins of *Salmonella* for the 3 conditions (Table S5); of these, 246 proteins were differentially expressed in response to H_2_O_2_ with *p* values ≤ 0.05 and 90% CI. Proteomics analysis showed that 121 and 125 proteins were upregulated and downregulated in response to stress by H_2_O_2_, respectively. Since Tn-seq revealed genetic requirements for fitness under the selection conditions, the identified genes are expected to express corresponding proteins under the conditions to perform their cellular functions. Often the proteins required for fitness under a given condition are overexpressed under the condition, but it may not be the case for some proteins. In this study, we had a unique opportunity to comparatively analyze both Tn-seq and the MS data to understand the relationship between genetic requirements and changes in expression level under the condition of interest, which was H_2_O_2_ in this study. We also obtained the list of essential genes based on our Tn-seq data, which could not tolerate insertions by definition, and if we were not certain about essentiality of a gene from our Tn-seq data, the gene was searched for essentiality in the previously reported list of *Salmonella* essential genes 22. The comprehensive list of essential genes allowed us to study any correlation between the essentiality and the changes in protein expression. Among the 246 proteins, there were 78 essential and 168 non-essential proteins. Among the 78 essential proteins, 25 were upregulated whereas 53 were downregulated. On the contrary, the majority (n = 96) of the detected non-essential proteins were upregulated, while 72 non-essential proteins were downregulated. To further examine the quantitative relationships closely, 64 genes/proteins identified by both methods (Table S5) were focused on. Among the 64 genes/proteins, 57 genes showed negative Log2FC based on Tn-seq data, and 41 proteins among the 57 were upregulated at protein level. However, only 12 proteins had *p* values of ≤ 0.05 (AhpC, ArcA, Crr, DksA, FliC, IcdA, OxyR, Pgm, RecA, RpoS, SlpA, and WecE).

Using KEGG pathway analysis, 150 proteins among the 246 were enriched in 21 pathways (Table S6). Interestingly, of the all 59 30S and 50S ribosomal proteins in *S.* Typhimurium, 37 of these proteins (63%) were downregulated in response to H_2_O_2_. Moreover, of the 8 identified proteins in TCA cycle, 6 proteins were downregulated, including 2 essential proteins.

Although DDA method can be used to search for all proteins in a complex sample, it is prone to miss identification of important proteins due to the fact that fragmentation of tryptic peptides from these proteins may not be triggered as a result of lower peptide ion intensities compared to the threshold set. To quantify proteins expressed for the genes identified by Tn-seq more precisely and accurately, we used targeted-proteomic approach by employing liquid chromatography coupled with triple quadrupole mass spectrometry (LC-QQQ-ESI-MS). Here, tryptic peptides of the protein were targeted for fragmentation (MS/MS) independent of their intensities, as described in Materials and Methods, and the observed sequence specific fragment ion intensities from three unique tryptic peptides were utilized for protein quantitation. Of the 137 Tn-seq identified genes, we selected 33 genes to quantify their proteins in response to H_2_O_2_ by using targeted proteomics (Table S5). Interestingly, 23 (70%) of the 33 tested proteins were upregulated in response to H_2_O_2_. This shows a good agreement between the results of the Tn-seq and the targeted proteomics.

### Aromatic amino acid biosynthesis genes are required for H_2_O_2_ resistance

Interestingly, our Tn-seq data revealed that the aromatic amino acid biosynthesis and metabolism pathway play a role in conferring resistance in *Salmonella* to H_2_O_2_ (Fig. 3A and 3B). Five genes, *aroB*, *aroD*, *aroE_2*, *aroK*, and *aroA* in the aromatic amino acid biosynthesis pathway were identified by Tn-seq, and the fitness of the mutants were significantly reduced in the presence of H_2_O_2_. To confirm this, 4 of these genes were evaluated using individual mutant assays. The *Salmonella aroK* mutant showed the strongest phenotype, because it failed to grow in the presence of H_2_O_2_L or H_2_O_2_H during 24 h incubation time. Also, the *aroB* mutant exhibited a strong phenotype, significantly extending lag phase for both H_2_O_2_ conditions. The *aroE_2* mutant also exhibited an extended lag time, but the *aroA* mutant did not show any difference in growth phenotype in the presence of H_2_O_2_. In addition, targeted-proteomics also showed that all these 5 proteins were upregulated in response to H_2_O_2_ (Fig. 3C and Table S5).

**Figure 3.**
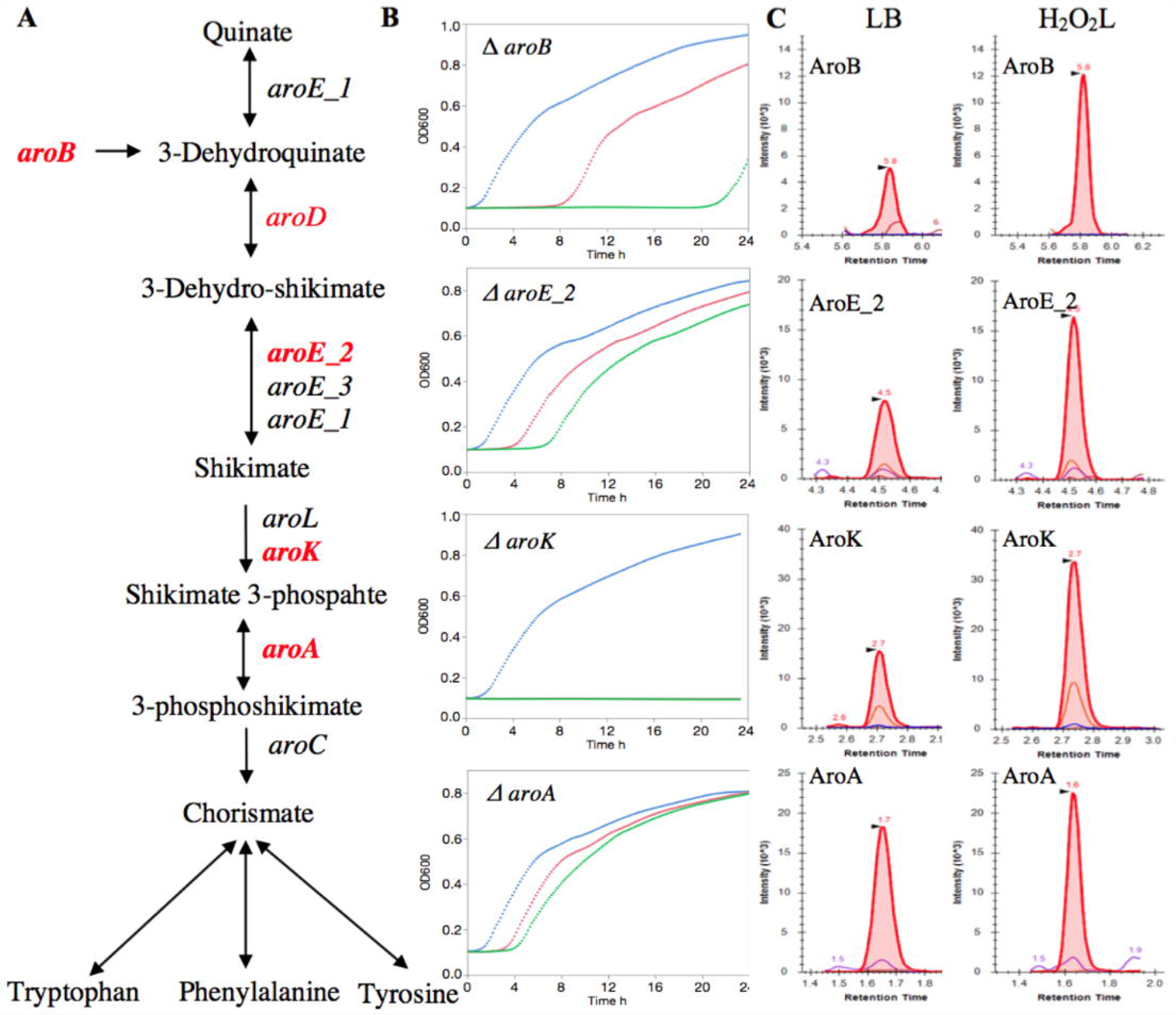
The role of aromatic amino acid biosynthesis genes in resistance to the H_2_O_2_. (A) Schematic representation of aromatic amino acid biosynthesis, adapted from the KEGG pathway database. The genes in red color were identified by the Tn-seq for H_2_O_2_ resistance in *Salmonella*. The red bold color genes were identified by the Tn-seq and the phenotypes were validated by the individual mutant assays. (B) The overnight cultures of the individual mutants were diluted 1:200 in the LB (no H_2_O_2_) and the LB contains either 2.5 mM H_2_O_2_ (H_2_O_2_L) or 3.5 mM H_2_O_2_ (H_2_O_2_H). The cultures were incubated at 37°C for 24 h in a 96-well plate. The colors of growth curve figures are blue for LB, red for H_2_O_2_L, and green for H_2_O_2_H. In the *aroK* growth curve, Δ the red color is under the green color. (C) Differential expression of *Salmonella* proteins in response to the H_2_O_2_L compared to the LB. Wild type *Salmonella* was grown in LB, H_2_O_2_L, and H_2_O_2_H until mid-exponential phase. Targeted-proteomics was quantified AroB, AroE_2, AroK, and AroA protein expressions in response to H_2_O_2_L. The shown peaks represent a unique peptide of the three peptides that were used of protein expression analysis.

ROS damages a variety of biomolecules via Fenton reaction, which consequently lead to metabolic defects, specifically auxotrophy for some aromatic amino acids 10. *E. coli* mutants that lack superoxide dismutase enzymes are unable to grow *in vitro* unless the medium are supplemented with aromatic (Phe, Trp, Tyr), branched-chain (Ile, Leu, Val), and sulfur-containing (Cys, Met) amino acids^36^. We identified the genes in the aromatic amino acid biosynthesis pathway that are critically important for resistance to H_2_O_2_. In this pathway, *aroK* catalyzes the production of shikimate 3-phosphate from shikimate, which consequently leads to the production of tryptophan, phenylalanine, tyrosine and some metabolites from the chorismate precursor in *E. coli*. Further, *aroK* mutant in *E*. coli displays increased susceptibility to protamine, a model cationic antimicrobial peptide. It has been suggested that resistance to protamine is probably due to the aromatic metabolites and product of *aroK* gene, which act as a signal molecule to simulate the CpxR/CpxA system and Mar regulators^37^. In our Tn-seq data, *cpxR*/*cpxA* and *marBCRT* were in the list of non-required genes, but the proteomics data indicated that CpxR was upregulated. Also, *aroK* mutant in *E. coli* is resistance to mecillinam, a beta-lactam antibiotic specific to penicillin-binding protein 2. It has been concluded that the AroK has a secondary activity in addition to the aromatic amino acid biosynthesis, probably related to cell division^38^. In addition, *aroK* gene presents a promising target to develop a non-toxic drug in *Mycobacterium tuberculosis* because *aroK* is the only *in vitro* essential gene among the aromatic amino acid pathway genes and blocking *aroK* kills the bacterium *in vivo*^39^.

Moreover, *aroK* gene plays a general role in *S.* Typhimurium persistence in pigs^40^. The *aroB* is another gene in the pathway that was identified by Tn-seq, which encodes 3-dehydroquinate synthase in the Shikimate pathway, aromatic amino acid biosynthesis pathway. *Salmonella* lacking *aroB* showed a strong growth defect in the presence of H_2_O_2_. When this mutant grown in the presence of H_2_O_2_, it increased the lag phase time by a 114-fold for the H_2_O_2_L and a 347-fold for the H_2_O_2_H as compared to the mutant grown in absence of H_2_O_2_. *S.* Typhimurium mutant lacking the *aroB* gene is attenuated in BALB/c mice^41^. In addition to *aroK* and *aroB*, *aroE_2* was also shown to be important for resistance to H_2_O_2_, because deletion of the *aroE_2* reduced the growth rate by 35% in the presence of H_2_O_2_ and increased the lag phase time, too. All these 3 genes in this pathway are required for systemic infection of *Salmonella* in BALB/c mice in a more recent study^18^. We observed that there was a strong correlation between the fitness based on Tn-seq data, growth rates measured by individual mutant assays, and upregulation of their proteins quantified via targeted proteomics. This demonstrates the power of proteogenomic approach in discovering and characterizing the genes that are required for growth under a specific condition.

### The *ybbM*, *ybbK*, and *ybbL* have a role in H_2_O_2_ resistance

The mutants with single deletion in each of *ybbK*, *ybbL*, and *ybbM* genes on the same pathway showed a strong phenotype against the activity of H_2_O_2_ in a dose-dependent manner. Based on Tn-seq data, the fitness of *ybbM* was −1.16 and −1.79 for H_2_O_2_L and H_2_O_2_H, respectively (Fig. 4A). The *ybbM* mutant demonstrated decreased growth rate by 38% for H_2_O_2_L and 100% for the H_2_O_2_H as compared to the mutant grown in the absence of H_2_O_2_. This mutant also increased the lag time by a 126-fold and a 267-fold for H_2_O_2_L and H_2_O_2_H, respectively (Fig. 4B). Also, the fitness score of *ybbK* was −0.92 for H_2_O_2_L and −1.81 for H_2_O_2_H. The *ybbK* mutant showed decrease of growth rate for H_2_O_2_L and H_2_O_2_H by 85% and 95%, respectively. The deletion increased the lag phase by a 46-fold and a 114-fold in the presence of H_2_O_2_L and H_2_O_2_H, respectively (Fig. 4B). Moreover, the fitness of *ybbL* mutant was −1.05 and −1.73 for H_2_O_2_L and H_2_O_2_H, respectively. Deleting the *ybbL* in *Salmonella* led to decrease in growth rate by 27% for H_2_O_2_L and 92% for H_2_O_2_H. The lag phase time for this mutant increased by a 22-fold and a 33-fold for H_2_O_2_L and H_2_O_2_H, respectively. In addition, YbbM, YbbL, and YbbK proteins were upregulated in response to H_2_O_2_; YbbM was the most upregulated protein among the 3 proteins (1.46-fold), followed by YbbL (1.29-fold), and YbbK (1.25-fold) (Fig. 4C and Table S5). The fitness scores of the Tn-seq of these 3 genes are correlated strongly with the growth rate, lag time of their respective mutants, and upregulation of their proteins. As the number of reads depletes after the selection for a mutant, (i) there was more reduction in growth rate, (ii) the mutant stays longer in the lag phase, and (iii) the protein expression elevates. These observations clearly point to their role in conferring resistance to the H_2_O_2_-mediated stress. These genes were described in the *Salmonella* reference genome as follows: *ybbM*, putative YbbM family transport protein, metal resistance protein; *ybbK*, putative inner membrane proteins; *ybbL*, putative ABC transporter, ATP-binding protein YbbL. To the best of our knowledge, there is only one published study on the *ybbM* and *ybbL*^42^. Based on their findings, YbbL and YbbM have a role in iron homeostasis in *E. coli* and are important for survival when the bacterium was challenged with 10 mM H_2_O_2_ for 30 min; this putative ABC transporter transports iron and lessens ROS species formation that generates via H_2_O_2_. In this study, we identified an additional gene, *ybbK*, in the same pathway as the gene required for resistance to H_2_O_2_, strongly establishing the role of these 3 genes in resistance to H_2_O_2_.

**Figure 4.**
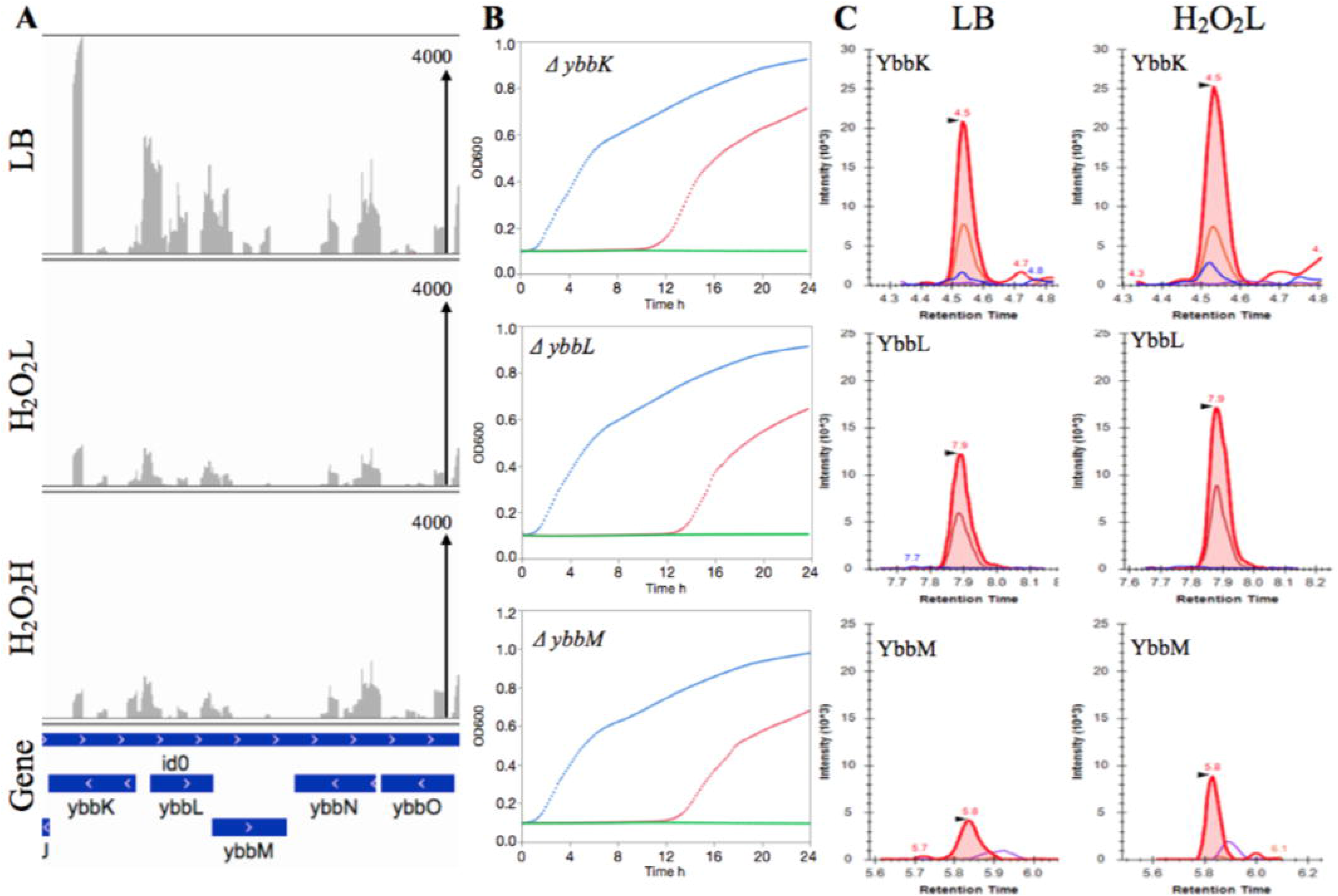
The *ybbK*, *ybbL*, and *ybbM* have a role in resistance to H_2_O_2_. (A) The Tn-seq profile of the LB (no H_2_O_2_), H_2_O_2_L (2.5 mM), and H_2_O_2_H (3.5 mM). It shows ∼6 kb of *Salmonella* Typhimurium genome starts with *ybbK* and ends with *ybbO*, horizontal axis. The read scale for the conditions are 4000, vertical axis. (B) The growth curve of *yybK*, Δ *yybL*, and Δ *yybM*. The overnight cultures of these three mutants were diluted 1:200 in LB, H_2_O_2_L, and Δ H_2_O_2_H. The cultures were incubated at 37°C for 24 h in a 96-well plate. The growth curve colors are blue which represents LB, red is H_2_O_2_L, and green is H_2_O_2_H. (C) Wild type *Salmonella* was grown in LB, H_2_O_2_L, and H_2_O_2_H until mid-exponential phase. Targeted-proteomics was quantified YbbK, YbbL, and YbbM protein expressions in response to H_2_O_2_L. The shown peaks represent a unique peptide of the three peptides that were used of protein expression analysis.

### The H_2_O_2_ scavenging and degrading genes

*Salmonella* employs redundant enzymes to degrade or scavenge ROS. The *katE*, *katG*, and *katN* genes encode catalases, which are involved in H_2_O_2_ degradation. The *ahpCF*, *tsaA*, and *tpx* genes encode peroxidases, which scavenge H_2_O_2_. The *sodA*, *sodB*, *sodCI*, and *sodCII* genes encode superoxide dismutases and these enzymes specifically scavenge O_2_^11, 12, 43-45^. However, none of these were present in the list of genes identified by Tn-Seq. Even though *katE*, *katG*, *ahpC*, *sodA*, *sodCI*, and *sodCII* showed reduced fitness, they did not meet the statistical threshold. However, the proteomics data indicated that AhpC (1.48-fold), SodB (1.46-fold), and TpX (1.39-fold) were upregulated in the presence of H_2_O_2_ (Table S5) and KatG was also upregulated, but its *p* value was 0.054. This reveals that these 4 proteins were the most important enzymes for H_2_O_2_ resistance under our experimental conditions. *Salmonella* containing an *ahpC* promoter-gfp fusion shows that expression of the *ahpC* is regulated by ROS that is generated from macrophages or exogenous H_2_O_2_ and the response to H_2_O_2_ is in a dose-dependent manner^46^.

*Salmonella* mutant that lacks *katE*, *katG*, or *ahpCF* can degrade micromolar concentrations of H_2_O_2_. However, *Salmonella* mutant that has deletions in the all 5 genes, *katE*, *katG*, *katN*, *ahpCF* and *tsaA* (HpxF), cannot degrade H_2_O_2_, is unable to proliferate in macrophages, and show reduced virulence in mice^11^. This emphasizes that *ahpC*, *sodB*, and *tpx* may be the primary players in scavenging and degrading H_2_O_2_ in our experiment. Why Tn-seq did not detect any of these genes, while proteomics detected only these 3 proteins among others? It may reflect the functional redundancy in the genetic network that prevented single deletions in one of these genes from exhibiting fitness defect. Alternatively, when these mutants were grown together with all other mutants in the library, the functional protein lacking in one mutant due to Tn5 insertion could have been compensated by the other mutants in the library.

In addition to these genes, *oxyR* was detected by Tn-seq (Fig. 1C) and DDA proteomics. The *oxyR* was identified by all 3 analysis methods of Tn-seq data and it was on the top of the list, indicating a severe fitness defect of the mutant. The *oxyR* gene encodes H_2_O_2_ sensor and transcription factor, which mediates protection against ROS. The *katG* and *ahpCF* are regulated by OxyR, peroxide response regulator^13, 14^. Although *Salmonella* OxyR regulon is induced in the *Salmonella*-containing vacuole in macrophage, the *oxyR* mutant was virulent in a BALB/c mouse and can grow well in human neutrophils *in vitro*^47, 48^. The fitness of *oxyR* mutant was reduced for both H_2_O_2_L and H_2_O_2_H with the respective fitness score of −4.96 and −5.94. *Salmonella oxyR* mutant exhibited a growth rate reduction by 24% and 40% for H_2_O_2_L and H_2_O_2_H, respectively. Comparing this reduction in growth rate to the other mutants such as *rpoS* or *aroK*, we observed that the *oxyR* mutant did not show severe phenotype and the mutant escaped from the lag phase easily. Moreover, our targeted proteomics indicated that the OxyR was not upregulated significantly. Further studies are needed to uncover the exact role of OxyR in response to ROS. However, previous studies implied that OxyR plays an essential role in resistance to H_2_O_2_ by regulating other proteins. OxyR induces Dps in *E. coli*, a ferritin-like protein that sequesters iron^49^. Sequestering of iron impairs the Fenton reaction, which consequently provides protection against ROS and reduces the damage of biomolecules. The *dps* gene was identified by the Tn-seq and its fitness score was −2.48. However, the Dps protein was downregulated based on the DDA proteomic analysis. To confirm this unexpected finding, we conducted the proteomic assay twice and each time with at least 4 technical replicates, but the Dps protein was significantly downregulated with *p* = 0.001. Further, the targeted-proteomics demonstrated the same result, pointing to the downregulation of Dps in response to H_2_O_2_. This is contrary to the previously reported works on Dps in *Salmonella* and the reason for the discrepancy is unclear.

### DNA repair system is important for H_2_O_2_ resistance

The imposed exogenous H_2_O_2_ activates DNA repair system in *Salmonella* in order to repair or eliminate the damage that occurred on the nucleotides. The *E. coli* RecA protein repairs double-strand DNA lesions through recombination^50^. In our Tn-seq analysis, the fitness score of this mutant was −5.36 for both concentrations, and in proteomics, the RecA was upregulated (1.79-fold). *Salmonella recA* mutant decreased the maximum OD_600_ by 16% for H_2_O_2_L and 22% for H_2_O_2_H as compared to the same mutant grown in LB. *Salmonella recA* mutant was also sensitive to exogenous H_2_O_2_ in aerated rich medium^26^. Moreover, *recG*, recombination and DNA repair gene^51^, showed a stronger phenotype than *recA* mutant. The *regG* deletion in *Salmonella* caused the growth rate reduction by 52% for H_2_O_2_L and 60% for H_2_O_2_H. This disruption in *recG* also caused the cells to stay in lag phase for a longer time in the presence of H_2_O_2_ as compared to LB; the lag time increased by a 62-fold and a 159-fold for H_2_O_2_L and H_2_O_2_H, respectively. In the blood of patients with *Salmonella* Typhi bacteremia, the proteins encoded by *recA*, *recG*, and *xthA* genes were detected, suggesting these proteins are actively expressed in the blood environment^52^. The XthA protein is another enzyme that participates in DNA repair mechanism induced by H_2_O_2_ and iron-mediated Fenton reaction. The *xthA* encodes exonuclease III, which repairs the damaged DNA. We found that the *xthA* gene was required based on the Tn-seq assay and its mutant had a reduced fitness score of −3.06 for H_2_O_2_L and a −4.38 for the H_2_O_2_H. Further, *Salmonella* lacking the *xthA* increased the lag time by 8-fold and a 12-fold for H_2_O_2_L and H_2_O_2_H, respectively. Targeted-proteomics showed upregulation of XthA (1.64-fold) in response to H_2_O_2_. *Salmonella* enterica serovar Enteritidis defective in *xthA* is susceptible to egg albumin^53^. *E. coli xthA* mutant is hypersensitive to H_2_O_2_. The *xthA* is also required for *Mycobacterium tuberculosis* to infect C57BL/6J mice^55^. In addition to the aforementioned genes involved in DNA repair system, *uvrA* encoding Holliday junction DNA helicase motor protein, *uvrC* encoding exonuclease ABC subunit A, *uvrD* encoding DNA-dependent helicase II, and *polA* encoding DNA polymerase I were among top of the list of the genes identified by Tn-seq as required for resistance to H_2_O_2_. Collectively, DNA repair system is crucial for the survival of the *Salmonella* in a niche that contains H_2_O_2_.

### Flagellar genes have a role for H_2_O_2_ resistance

Some flagellar genes, *fliC* and *fliB*, were shown to be important for resistance to H_2_O_2_. These two genes were identified by Tn-seq and their proteins were shown to be upregulated. *Salmonella* lacking either of these genes exhibited a strong phenotype in the presence of H_2_O_2_. During 24 h of incubation, *fliC* and *fliB* mutants could not grow in both H_2_O_2_ conditions. However, growth of *fliD* mutant was not affected by H_2_O_2_. The *fliC* was shown to have a role in *Salmonella* Typhi interaction with human macrophages and *Salmonella* Typhimurium *fliB* mutant was defective in swarming motility^56, 57^. Currently it remains unclear how flagella genes can be involved in the resistance of *Salmonella* to oxidative stress, which warrants future study into this direction.

### Fe-S cluster biogenesis system is required for H_2_O_2_ resistance

*Salmonella* requires the genes from Fe-S cluster biogenesis system in order to resist H_2_O_2_. Our Tn-seq analysis identified 5 genes in this system that are required for the resistance. In *isc* operon (Fe-S cluster), *iscA*, *hscB*, and *hscA* were among the genes required to resist H_2_O_2_. Particularly, the *hscA* is on the top of the gene list identified by Tn-seq. In *E. coli*, this operon is regulated by *iscR*, iron sulfur cluster regulator^58^; in *Salmonella* the gene *iscR* encoding this transcription regulator is named *yfhP*. The HscB and HscA chaperones are believed to be involved in the maturation of Fe–S proteins^59, 60^. The second operon that is involved in Fe-S protein biogenesis is the *suf*, sulfur mobilization operon. Tn-seq found that two genes in this operon were required for *Salmonella* to resist H_2_O_2_; *sufS* and *sufC*. *Salmonella* mutant lacking *sufS* exhibited a strong phenotype in the presence of H_2_O_2_ and could not grow during the 24 h of incubation as compared to LB. The SufS with SufE in *E. coli* form a heterodimeric cysteine desulphurase and SufB, SufC, and SufD form a pseudo-ABC-transporter that could act as a scaffold^60^; this operon is regulated by OxyR^14^. The other known genes in these two operons that are present in *Salmonella* are *iscA*, *sufA*, *sufB*, and *sufD*; they showed a reduced fitness, while their *p* values were > 0.05. The damage of Fe-S clusters is not only problem for the defective proteins, but also it fuels the Fenton reaction via the released iron and H_2_O_2_^10^. Thus, *Salmonella* uses Fe-S cluster repair system as an arsenal to overcome the damage imposed by H_2_O_2_.

### DNA adenine methylase is important for H_2_O_2_ resistance

DNA adenine methylase genes, *dam* and *damX*, are involved in *Salmonella* resistance against H_2_O_2_. Our Tn-seq data showed that fitness of *dam* and *damX* mutants were reduced in the presence of H_2_O_2_. To confirm this, *Salmonella dam* mutant was grown in both conditions. Under H_2_O_2_L, the growth rate was reduced by 42% as compared to the mutant in LB and the mutant could not grow under H_2_O_2_H during the 24 h of incubation. In addition, the lag time of the *Salmonella dam* mutant was extended by a 19-fold for H_2_O_2_L. While the *Salmonella damX* mutant displayed a moderate phenotype as compared to the *dam* mutant, the *damX* mutant also showed that the growth rate decreased by 23% and 33% for H_2_O_2_L and H_2_O_2_H, respectively. The lag time was extended for this mutant by a 25-fold and a 49-fold for H_2_O_2_L and H_2_O_2_H, respectively. The *dam* regulates virulence gene expression in *S*. Typhimurium^61^. The different levels of Dam affects virulence gene expression, motility, flagellar synthesis, and bile resistance in the pathogenic *S*. Typhimurium 14028S^62^. Dam methylation activates the gene that are involved in lipopolysaccharide synthesis^63^. Moreover, *Salmonella* defective in *damX* is very sensitive to bile^64^. Collectively, our study demonstrates the critical role of DNA adenine methylase in *Salmonella* resistance against H_2_O_2_.

### Other genes for H_2_O_2_ resistance

Beside the important pathways described above, there were many additional genes also important for resistance to H_2_O_2_. Among those, the 3 unrelated genes, *rpoS*, *pgm*, and *tonB*, are important ones that deserve more attention. The *rpoS* mutant showed reduced fitness and its protein was upregulated in the presence of H_2_O_2_. *Salmonella* mutant defective in *rpoS* showed a strong phenotype when grown in the presence of H_2_O_2_. The *rpoS* encodes the alternative sigma _S 65_ factor σ^S^, subunit of RNA polymerase; it is the master regulator of stress response^65^. This implies that *rpoS* is an important component of the genetic regulatory network that *Salmonella* employs in order to resist H_2_O_2_. Furthermore, the fitness of *pgm* mutant was reduced and its protein was upregulated in the presence of H_2_O_2_. Knock out of *pgm* in *Salmonella* caused a decrease in growth rate, increase the lag phase time, and reduce the maximum OD_600_ in the presence of H_2_O_2_. The *pgm* encodes phosphoglucomutase which required for catalysis of the interconversion of glucose 1-phosphate and glucose 6-phosphate^66^. This gene contributes to resistance against antimicrobial peptides, is required for *in vivo* fitness in the mouse model, and participates in LPS biosynthesis^67^. Lastly, the fitness of *tonB* mutant was also reduced. *Salmonella* lacking *tonB* exhibited a strong phenotype in the presence of H_2_O_2_ as compared to the mutant grown in LB. The gene mediates iron uptake in the *Salmonella*^45^. In addition, seven of the genes identified in our study (*proC*, *arcA*, *barA*, *exbD*, *flhD*, *fliC*, and *fliD*) were previously shown to be important for interaction of *Salmonella* Typhi with human macrophages^56^.

In summary, we applied Tn-seq and proteomic analysis to find the genes and proteins that are required in *S.* Typhimurium to resist H_2_O_2_ *in vitro*. As the concentration of H_2_O_2_ increased, the growth rate reduced, the lag time extended, the fitness of mutants decreased, and some proteins were differentially expressed. Validation of Tn-seq results with individual mutant assays indicated the accuracy of the identified genes in response to the two H_2_O_2_ concentrations. The targeted-proteomics had a good agreement with Tn-seq. We found many genes that have not been associated to resistance to H_2_O_2_ previously and these genes will be focus of our future research. *Salmonella* employs multiple pathways to resist H_2_O_2_ and the most important ones are ROS detoxifying enzymes, amino acid biosynthesis (*aroK* and *aroB*), putative iron transporters (*ybbK*, *ybbL*, *ybbM*), iron homeostasis, Fe-S cluster repair, DNA repair, flagellar and DNA adenine methylase genes. The genes identified in this study will broaden our understanding on the mechanisms used by *Salmonella* to survive and persist against ROS in macrophages.

Our unbiased system-wide approach, Tn-seq, was successful in identifying novel genetic determinants that have not been implicated previously in *Salmonella* resistance to oxidative stress. Furthermore, the combined use of quantitative proteomic approach has provided additional insights on the function or mode of action of the identified genetic determinants in resisting oxidative stress. As expected, the majority of the proteins important for resistance to H_2_O_2_ were upregulated in response to the same stressor. However, the expression level did not increase for some proteins, in spite of their known roles in resistance to H_2_O_2_. Interestingly, the downregulation of Dps and other proteins was counterintuitive to the common mode of protein regulation and function, yet it may point to some unknown aspects of how *Salmonella* regulates the expression of those proteins to better cope with the oxidative stress during infection in macrophage. The genes identified in this study will broaden our understanding on the mechanisms used by *Salmonella* to survive and persist against ROS in macrophages.

## Methods

### Construction of Tn5 mutant library

We mutagenized *Salmonella* enterica subsp. enterica serovar Typhimurium str. ATCC 14028S (with spontaneous mutation conferring resistance to nalidixic acid (NA)), by biparental mating using *Escherichia coli* SM10 λ*pir* carrying a transposon-delivery plasmid vector pBAM1 λ (Ampicillin (Amp) resistance) as a donor strain^68^. The plasmid pBAM1 was generously provided by Victor de Lorenzo (Systems and Synthetic Biology Program, Centro Nacional de Biotecnología, Madrid, Spain). The donor strain*, E. coli* Sm10 λpir (pBAM1), was grown overnight in LB with 50 µ g/ml Amp and recipient strain was grown in LB with 50 µ g/ml NA at 37°C. Equal volumes (1 ml) of donor and recipient were mixed, centrifuged, washed in 10 mM MgSO_4_, and re-suspended in 2 ml PBS (pH 7.4). Then, the mating mixture was concentrated and laid on a 0.45-µ m nitrocellulose filters (Millipore). The filter was incubated at 37°C on the surface of LB agar plate. After 5 h of conjugation, the cells on the filter was washed in 10 mM

MgSO_4_, and resuspended in 1 ml MgSO4. The conjugation mixture was plated on LB agar containing 50 µ g/ml NA and 50 µ g/ml kanamycin (Km). After approximately 24 h at 37°C, we scraped the colonies into LB broth containing 50 µ g/ml Km and 7% DMSO. The yield was approximately 68,000 individual colonies from each conjugation. Five independent conjugations were conducted to yield approximately 325,000 mutants. The library was stored at −80°C in aliquots. To determine the frequency of the mutants that have been produced by integration of the entire delivery plasmid, the colonies were picked randomly and streaked on LB plates (Km) and LB plates (Km and Amp). It was shown that ∼20% of the cells in the library were resistant to Amp, indicating a significant portion of the Km-resistant colonies was not from authentic transposition events.

### Measuring growth responses of *S.* Typhimurium to H_2_O_2_

To determine the effect of H_2_O_2_ concentrations on growth parameters, overnight culture of the wild type *S.* Typhimurium 14028s was inoculated into fresh LB broth media (1:200 dilution) to give a seeding concentration corresponding to OD_600_ of ∼0.1. The LB broth contained freshly prepared H_2_O_2_ to give the final concentrations ranging from 0.05 to 10 mM. The cultures were directly added into 96-well microplate (200 μl/well). The microplate was incubated in a Tecan Infinite 200 microplate reader at 37°C, with shaking duration 5 s, shaking amplitude 1.5 mm, and reading OD_600_ every 10 min. The number of replicates were at least three. The lag time, growth rate, and maximum OD600 were calculated using GrowthRates script^69^. Growth Rate % decrease was calculated as follows^70^: Growth Rate % decrease = ((μPC – μ_S_)/μPC) × 100; where μ = the maximum slope (growth rate), μ_PC_ = growth rate of positive control (without H_2_O_2_), μ_S_ = growth rate in the presence of H_2_O_2_.

### Selection of the mutant library for Tn-seq analysis

The transposon library was thawed at room temperature and diluted 10^−1^ in fresh LB broth. To activate the library, the diluted library was incubated at 37°C with shaking at 225 rpm for an hour. Then, the culture was washed twice with PBS and resuspended in LB broth medium. The library was inoculated to 20 ml LB broth and LB broth supplemented with either 2.5 or 3.5 mM H_2_O_2_ (H_2_O_2_L and H_2_O_2_H, respectively), seeding CFU was 3.5×10^6^ per ml. Then, when the cultures reached mid-exponential phase, OD600 of 2.7 (∼1.17×10^8^ CFU/ml), the incubation was stopped, and the culture was immediately harvested by centrifugation, and stored at −20°C.

### Preparation of Tn-seq amplicon libraries

Genomic DNA was extracted from the harvested cells using DNeasy Blood & Tissue kit (Qiagen), and quantified using Qubit dsDNA RB Assay kit (Invitrogen). As described above, 20% of the mutants in the library were the result of the integration of pBAM1 into chromosome. To remove the Tn5-junction sequences originated from the plasmid in the Tn-seq amplicon libraries, genomic DNA was digested with PvuII-HF (New England Biolabs), which digests immediately outside the inverted repeats on both sides of Tn5 in pBAM1, and purified with DNA Clean & Concentrator-5 kit (Zymo Reaerch). Then, a linear PCR extension was performed using a Tn5-specific primer in order to produce single stranded DNA corresponding to Tn5-junction sequences. To increase the specificity in extending into Tn5-junction sequences, the linear PCR was conducted with a dual priming oligonucleotide Tn5-DPO (5’-AAGCTTGCATGCCTGCAGGTIIIIICTAGAGGATC-3’) that is specific to Tn5 end^71^. The PCR reaction contained 25 μ M Tn5-DPO primer, 100 ng gDNA, and 50 μ MQ-H_2_O. The PCR cycle consisted of the initial denaturation at 95°C for 2 min, followed by 50 cycles at 95°C for 30 sec, 62°C for 45 sec, and 72°C for 10 sec. The PCR product was purified with DNA Clean & Concentrator-5 kit and eluted in 13 μ TE buffer. After that, C-tail was added to the 3’ end of the single-stranded DNA. The C-tailing reaction was consisted of 2 μ mM dCTP, 1 μ 1 mM ddCTP, 0.5 μ TdT and 13 μ μl CoCl_2_, 2.4 l 10 purified linear PCR product. The reaction was performed at 37°C for 1 h and the enzyme was inactivated by incubation at 70°C for 10 min. The C-tailed product was purified with DNA Clean & Concentrator-5 kit and eluted in 12 μ TE. Next, the exponential PCR was performed with forward primer, P5-BRX-TN5-MEO, 5’-AATGATACGGCGACCACCGAGATCTACACTCTTTCCCTACACGACGCTCTTCCGATCTNNNNAG-6 nt barcode-CCTAGGCGGCCTTAATTAAAGATGTGTATAAGAG-3’ and reverse primer, P7-16G, 5’-CAAGCAGAAGACGGCATACGAGCTCTTCCGATCTGGGGGGGGGGGGGGGG-3’ to attach Illumina adapter sequences along with the sample barcodes. The PCR reaction contained 25 μ μ M P5-BRX-TN5-MEO primer, 10 μM P7-16G primer, 1 μ l; the PCR condition started with initial denaturation at 95°C for 2 min, followed by 36 cycles of 95°C for 30 sec, 60°C for 30 sec, and 72°C for 20 sec, with the final extension at 72°C for 5 min. Then, the size selection of the DNA was performed using agarose gel electrophoresis. The 50 μl PCR products were incubated μ at 60°C for 15 min and incubated on ice for 5 min, and immediately loaded on the 1% agarose gel in 0.5% TAE buffer. After running the gel, the DNA fragment of size 325-625 bp was cut and put in a microtube for each sample. The DNA was extracted from the gel using Zymoclean Gel DNA Recovery kit (Zymo Reaerch). The prepared DNA libraries were quantified using Qubit dsDNA RB Assay kit. Since each library has its own barcode, the libraries were combined and sequenced on a flow cell of HiSeq 3000 using single end read and 151 cycles (Illumina) at the Center for Genome Research & Biocomputing in Oregon State University.

### Tn-seq data analysis

The preliminary data analysis was conducted by using a super computer in the High Performance Computing Center (AHPCC) at the University of Arkansas. The libraries that were multiplexed for sequencing were de-multiplexed using a custom Python script. The script searched for the six-nucleotide barcode for each library for perfect matches. In order to extract the transposon genomic junctions, we used Tn-Seq Pre-Processor (TPP) tool^30^ with some modifications in the script. The TPP searched for the 19 nucleotide inverted repeat (IR) sequence and identified five nucleotides (GACAG) at the end of the IR sequence, allowing one nucleotide mismatch. The Tn5-junctions that start immediately after GACAG were extracted and the C-tails at the end of junctions were removed. Tn5-junction sequences less than 20 nucleotides were discarded and remaining Tn5-junctions were mapped to the *Salmonella* enterica serovar Typhimurium 14028S genome and plasmid using BWA-0.7.12^72^. To identify genes that are required for H_2_O_2_ resistance, the following three Tn-seq analysis tools were used for comparative analysis: (i) ARTIST^28^: the genomic junctions were mapped to the reference genome using Bowtie 2.2.7^73^. The number of insertions and reads were determined for genes and intergenic regions. The data were normalized with default script in the ARTIST. Then, the relative abundance of Tn5 insertions in the output library versus the input were calculated. Later, the *p* values were calculated from a 100 independent Mann–Whitney U test (MWU) analysis that were carried out on input and output data for each gene. Finally, the genes were considered conditionally essential if the *p* values were ≤ 0.05 in the 90 of the 100 MWU tests. (ii) Tn-seq Explorer^29^: The output SAM files from the TTP were used as input to the Tn-seq Explorer. The unique insertions with less than 20 reads were removed from the input and outputs. Using the window size of 550 and excluding 5% of beginning of genes and 20% of the end of genes, Essentiality Index (EI), number of unique insertions, and total number of reads per gene were counted. The EI of more than 10 were removed from the input. Genes with less than 300 nucleotides were removed. Deferential EI were calculated from input and outputs (IE = output EI–input EI) and genes with Δ ΔIE more than −1 were removed. Log2 fold change of reads were calculated from input and output (Log2FC = log2 (output reads/input reads)) and the genes were ranked based on the Log2FC value from least value to highest. The genes with Log2FC value of less than −2 from the H_2_O_2_H and present in H_2_O_2_L were considered conditionally essential. (iii) TRANSIT^30^: The output wig files from the TTP was used as input data file for TRANSIT. The comparative analysis was conducted with Tn5 resampling option. The reads were normalized with trimmed total reads (TTR). Insertions outside the 5% and 10% sequences from 5’- and 3’- ends were removed, respectively. The genes were considered conditionally essential if *p* values ≤ 0.05.

### Phenotypic evaluation of individual deletion mutants

The mutants were obtained from *Salmonella enterica* subsp. *enterica*, 14028s (Serovar Typhimurium) Single-Gene Deletion Mutant Library through BEI Resources (www.beiresources.org). The overnight cultures of *S.* Typhimurium mutants were added into fresh LB broth media containing freshly prepared H_2_O_2_ (2.5, or 3.5 mM/ml) (1:200 dilution) to give seeding OD_600_ of 0.1. The cultures were directly added into 96-well microplates and incubated in Tecan Infinite 200 at 37°C for 24 h. The lag time, growth rate, and maximum OD_600_ were calculated using GrowthRates^69^.

### Sample preparation for proteomics and mass spectrometry analysis

The overnight culture of the wild type *S.* Typhimurium 14028 was diluted 1:200 in 50 ml LB medium, and LB containing either 2.5 or 3.5 mM H_2_O_2_ in a 300-ml flask. The cultures were grown until mid-exponential phase (OD_600_ of 2.7), and the 50 ml volume of cultures were used for a total protein extraction by using Qproteome Bacterial Protein Prep kit (Qiagen). Disulfide bonds within proteins were reduced with 2-Mercaptoethanol and separated by SDS-PAGE gel electrophoresis. For each condition, there were three lanes with approximately 300 μg of proteins. The gel then was stained via coomassie blue dye. The gel portions of 3 lanes for each condition were cut out and chopped into small pieces, pooled together, washed twice with 50 mM NH_4_HCO_3_, destained with NH_4_HCO_3_/ 50% Acetonitrile (ACN), and dried with pure ACN. Then, the proteins were reduced using 10 mM Dithiothreitol in 50 mM NH_4_CO_3_ and the alkylation was conducted with 10 mg/ml Iodoacetamide Acid in 50 mM NH_4_CO_3_. After that, the proteins were washed with NH_4_HCO_3_, and dried with pure ACN. Mass spectrometry grade Trypsin gold from Promega (∼20 ng/μl in 50 mM NH_4_HCO_3_) was added to dried gels, and left it overnight for efficient in-gel digestion of the proteins at 37°C. During the digestion, tryptic peptides diffused out into the solution. Gel pieces then were extracted three times by 50% CAN/0.1% TFA solution and incubated at 37°C for 15 minutes. Later, these digests were analyzed by ESI-LC-MS/MS at State Wide Mass Spectrometry Facility, University of Arkansas at Fayetteville. Data dependent analysis (DDA) for the in-gel trypsin digested samples from each condition were performed by using an Agilent 1200 series micro flow HPLC in line with Bruker Amazon-SL quadrupole ion trap ESI mass spectrometer (QIT-ESI-MS). All the ESI-MS analyses were performed in a positive ion mode using Bruker captive electrospray source with a dry nitrogen gas temperature of 200°C, with nitrogen flow rate of 3 L/minute. LC-MS/MS data were acquired in the Auto MS(n) mode with optimized trapping condition for the ions at m/z 1000. MS scans were performed in the enhanced scanning mode (8100 m/z/second), while the collision-induced dissociation or the MS/MS fragmentation scans were performed automatically for top ten precursor ions with a set threshold for one minute in the UltraScan mode (32,500 m/z/second). Tryptic peptides were separated by reverse-phase high-performance liquid chromatography (RP-HPLC) using a Zorbax SB C18 column, (150 × 0.3 mm, 3.5 µ m particle size, 300 Å pore size, Agilent Technologies), with a solvent flow rate of 4 µ L/minute, and a gradient of 5%–38% consisting of 0.1% FA (solvent A) and ACN (solvent B) over a time period of 320 minutes. Tryptic peptides were then identified by searching MS/MS data in *S.* Typhimurium 14028S reference proteome database^24, 35^ by using MASCOT database search software^34^. MS1 intensities of the integrated areas of these identified tryptic peptides were compiled and grouped in skyline software according to the replicates/conditions to perform statistical analysis. Targeted protein work were performed using Shimadzu UPLC-20A coupled to 8050 triple quadrupole ESI-MS with heated probe. Sequence specific fragment ion intensities from at least three unique tryptic peptides from the protein of interest were used in the protein quantitation. Multiple reaction monitoring (MRM) events corresponding to sequence specific fragment ions derived from the precursor tryptic peptides were targeted to operate at a certain specific retention time intervals predicted by in house retention time library. This library was generated using the correlation of relative hydrophobicity of the tryptic peptides with their retention times (RT) from highly common housekeeping proteins, for the UPLC method used in this analysis as described below. While the RT were correlated well within 99% confidence, sufficient number of sequence specific fragment ions were used as basis for identification of the tryptic peptide by MS/MS alone. Specificity and the confidence was achieved by incorporating RT prediction. In addition to the application of skyline in quantitation, skyline software was also used in predicting RT and optimizing parameters such as collision energies and voltages with the help of Shimadzu Labsolution software. Tryptic peptides were separated by reverse-phase ultra-high-performance liquid chromatography (RP-UPLC) compatible Shimadzu C18, 1.9-micron particle size, 50×2.1 mm column (SN # 16041880T), with a solvent flow rate of 0.3 mL/minute, and a gradient of 5%–90% consisting of 0.1% FA (solvent A) and 0.1% FA in ACN (solvent B) over a time period of 10 minutes.

### Accession numbers

Tn-seq sequencing data are available on NCBI Sequence Read Archive. LB: PRJNA352537; H_2_O_2_L: PRJNA352862; H_2_O_2_H: PRJNA352865.

## Acknowledgment

We thank Jack Jiang, a high school student, for writing the demultiplexing Python script. We are thankful for Dr. Thomas R. Ioerger and Dr. Michael A. DeJesus at Texas A&M University for the help on the use of TRNASIT. We extend our special thanks to Dr. Jeff F. Pummill and Dr. Pawel Wolinski at Arkansas High Performance Computing Center for their help to use the facility.

## Funding

This work was partially supported by Arkansas Biosciences Institute. The first author was supported by his parents, Cell and Molecular Biology (CEMB) program at the University of Arkansas, and Human Capacity Development Program-Kurdistan Regional Government (HCDP- KRG).

## Author Contributions

Conceived and designed the experiments: YK SK. Performed the experiments, analyzed the data, wrote the manuscript: SK. Proteomics work: RL SK AQ JL. Revised the manuscript: YK SK. All authors read the final version of the manuscript.

## Competing financial interests

The authors declare no competing financial interests.

## Supplementary information

**Figure S1. Stability of H_2_O_2_ in LB medium during the experiments**. LB broth media supplemented with freshly diluted 3.5 mM H_2_O_2_ at each of indicated time points. At 0 h, immediately after adding H_2_O_2_ to LB broth, the media inoculated with *Salmonella* Typhimurium. At 24 h, 24 hours before the inoculation media with bacteria H_2_O_2_ added to media, and at 11 d, 11 days before the inoculation media with bacteria H_2_O_2_ added. The media supplemented with H_2_O_2_ left at room temperature. LB was free of H_2_O_2_. The cultures were incubated at 37°C for 24 h in a 96-well plate with OD_600_ reading every 10 minutes.

**Figure S2. Tn-seq library preparation diagram for Illumina sequencing**. The genomic DNA extracted from the selected library and subjected to two PCR amplifications. First PCR was linear and specific forward primer used to capture and amplify Tn5 junctions. Second PCR was exponential and Illumina adaptors with a barcode added. The PCR product gel purified and sequenced on an Illumina platform.

**Figure S3. The reproducibility of Tn-seq.** Correlation between reads per ORFs of *Salmonella* Typhimurium Tn-seq conditions, H_2_O_2_L (2.5 mM) and H_2_O_2_H (3.5 mM). Two ORFs excluded in this correlation, STM14_2422 and STM14_2428.

**Table S1. Salmonella Typhimurium Tn-seq sequencing in numbers.** Number of extracted reads, mapped reads, and unique insertions for LB (H_2_O_2_ free), H_2_O_2_L (2.5 mM), and H_2_O_2_H (3.5 mM) presented. Mean length of mapped genomic junctions and hits per nucleotide shown.

**Table S2. Full list of Salmonella Typhimurium Tn-seq genome of the study.** The list of 137 *Salmonella* required genes for H_2_O_2_ resistance. The full data set of Tn-seq genome analysis with three tools, ARTIST, Tn-Seq Explorer, and TRANSIT for H_2_O_2_L (2.5 mM) and H_2_O_2_H (3.5 mM).

**Table S3. List of functional categories required for Salmonella Typhimurium H_2_O_2_ resistance.** SP_PIR_Keywords used with default options for functional categories analysis of the 137 genes that were required for H_2_O_2_ resistance in *S*. Typhimurium. The gene recognition by the analysis tool based on official gene symbols.

**Table S4. Lag time, growth rate, and maximum OD_600_ of Salmonella Typhimurium mutants.** 50 mutants and wild-type of *S*. Typhimurium grown in LB (H_2_O_2_ free), H_2_O_2_L (2.5 mM), H_2_O_2_H (3.5 mM). The cultures were incubated at 37°C for 24 h in a 96-well plate with OD_600_ reading every 10 minutes. Lag time, growth rate, and maximum OD600 calculated for each mutant and compared to the wild-type.

**Table S5. Full list of *Salmonella* Typhimurium proteomic analysis in response to H_2_O_2_.** *S*. Typhimurium strain 14028S grown in LB (H_2_O_2_ free), H_2_O_2_L (2.5 mM), H_2_O_2_H (3.5 mM) till mid-log phase. Proteome profiles analyzed by utilizing ESI-LC-MS/MS in data-dependent acquisition (DDA) mode and LC-QQQ-ESI-MS for targeted proteomics.

**Table S6. Deferentially expressed proteins of Salmonella Typhimurium in response to H_2_O_2_ and their pathways.** *S*. Typhimurium strain 14028S grown in LB (H_2_O_2_ free), H_2_O_2_L (2.5 mM), H_2_O_2_H (3.5 mM) till mid-log phase. KEGG pathway analysis used to categorize deferentially expressed proteins (p < 0.05). Blue for downregulated proteins, red for upregulated proteins and bold represents essential proteins.

